# Repressive *S*-adenosylmethionine biosynthesis status inhibits transcription of *HeT-A* retrotransposon in the germline of *Drosophila*

**DOI:** 10.1101/2025.04.12.648310

**Authors:** Yoshiki Hayashi, Shinjiro Hino, Tetsuya Sato, Soshiro Kashio, Kuniaki Saito, Ban Sato, Natsuko Kawano, Daisuke Saito, Masayuki Miura, Mikita Suyama, Mitsuyoshi Nakao, Satoru Kobayashi

## Abstract

*S*-adenosylmethionine (SAM) is the principal cellular donor of methyl moiety in the methylation reaction and regulates gene expression by regulating methylation-related cellular events, such as epigenetic status. Although SAM biosynthesis affects a variety of biological phenomena including disease and aging, whether cell-specific SAM biosynthesis status is present and how it contributes to cellular function are largely unknown. Here, we firstly showed that the *Drosophila* germline in gametogenesis has a repressive SAM biosynthesis status through the observation of SAM synthetase (Sam-S), a key enzyme for SAM biosynthesis. In addition, our study showed that germline-unique repressive SAM biosynthesis status contributes to inhibition of retrotransposon expression; enhancement of SAM biosynthesis in germline caused excessive expression of retrotransposons including *HeT-A*, a telomere-specific retroelement, as the most affected target. We found that the promoter activity of *HeT-A* is enhanced in SAM increased condition with increased accumulation of 6mA DNA methylation, the major DNA methylation modification in the *Drosophila* genome. Interestingly, the enhanced 6mA enrichment and gene expression in enriched loci were not correlated in other retrotransposons or structural genes. Taken together, our results suggest that SAM-deficient status in the germline uniquely regulates *HeT-A* transcription via 6mA methylation modification. Thus, our study provides a new understanding of how germline unique metabolic status contributes to regulation of the retrotransposon.

## Introduction

A growing body of evidence emphasizes that the cellular metabolisms are the one of the essential regulatory layers for gene expression by regulating epigenetic status (1). *S*-adenosylmethionine (SAM), which is synthesized from methionine (Met), is the essential donor of proteins or nucleic acids methylation modification to regulate a variety of biological processes, including cell differentiation, diseases, and aging (2–5). SAM synthesis depends on the evolutionally conserved enzyme SAM Synthase (Sam-S) (Figure 1A). After the synthesis, SAM is catalyzed by a variety of methyltransferases to be *S*-adenosylhomocysteine (SAH). Since SAM is the essential metabolite for methylation modification, and methylation modification occurs in basically all types of cells, the cell-type-specific status of SAM biosynthesis has not been elucidated.

**Figure 1.**
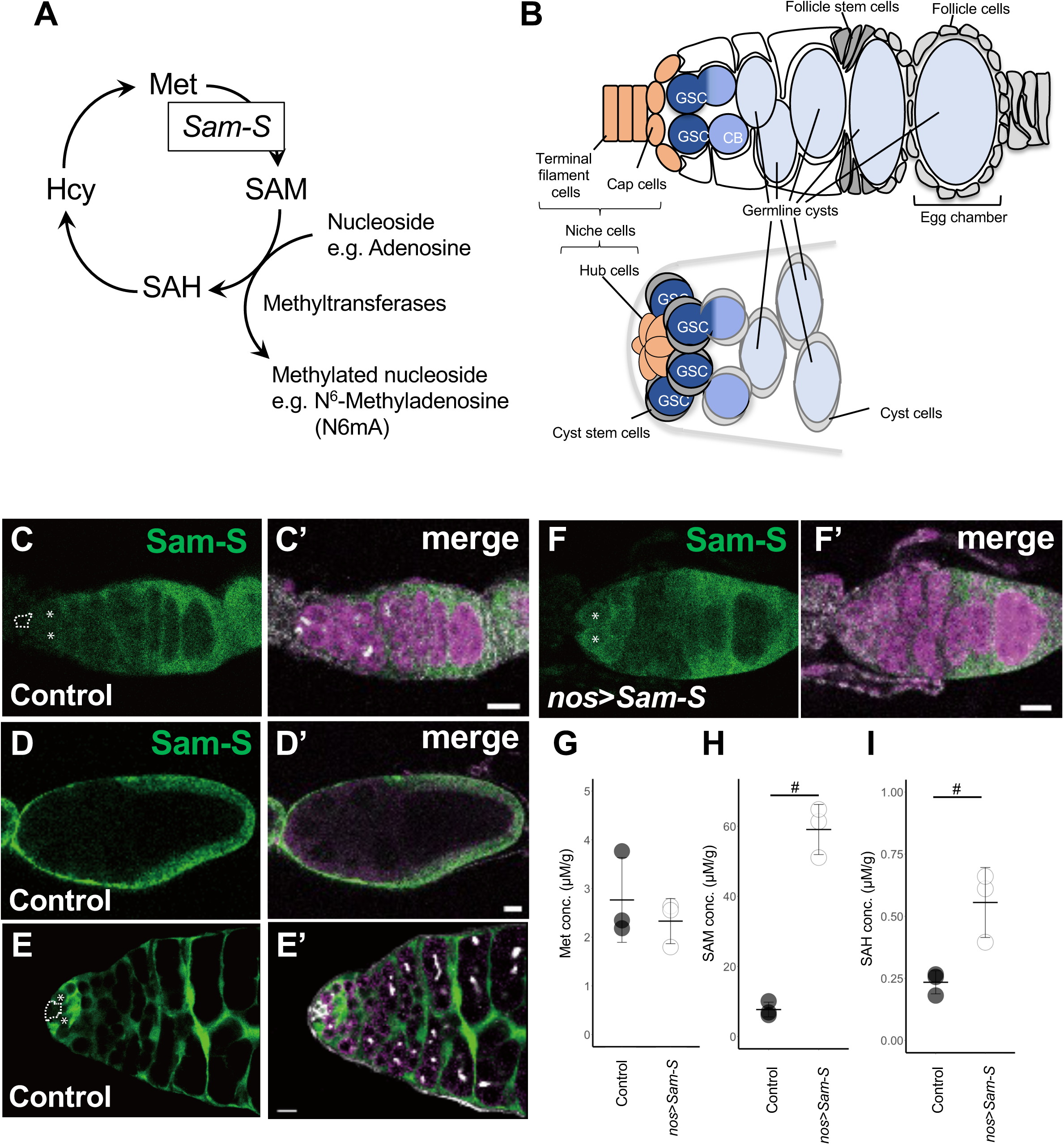
SAM biosynthesis status during *Drosophila* gametogenesis. (A) Schematic diagram of SAM biosynthesis. (B) Schematic diagram of early stage of *Drosophila* oogenesis (top) and spermatogenesis (bottom). (C, C’) Expression of Sam-S protein (Green) in early oogenesis in control females. The magenta signal in C’ indicates the germline, and asterisks indicate GSCs. Niche cells are marked with a dashed line in C. White signals in C’ indicate the signal from the 1B1 antibody, which marks the germline-specific organelle, the fusome. (D, D’) Expression of Sam-S protein (Green) in late oogenesis in wild type females. (E, E’) Expression of Sam-S protein (Green) in early spermatogenesis. The magenta signal indicates the germline, and the white signals indicate the signal from the 1B1 antibody and the FasIII antibody, which mark fusome and niche cells, respectively. Asterisks indicate GSCs, and a dashed line indicates niche cells (F, F’) Expression of Sam-S protein (green) in early oogenesis in *nos* > *Sam-S* females. Sam-S protein is detected in germline (magenta in F’) including GSCs (asterisks in F). (G-I) The measurements of Met (G), SAM (H) and SAH (I) amount in control (gray) and *nos* > *Sam-S* (white) ovaries. #: p < 0.05 (Student’s t-test). Bars in C-F’: 10µm.

*Drosophila* gonad provides an ideal model to investigate the presence and role of cell-type-specific SAM biosynthesis status, since it contains a variety of cell-types given rise from two fundamentally different cell lineages, germline and somatic line (Figure 1B). Although a previous study showed that SAM biosynthesis has a function to regulate germline aging during female gametogenesis (3), whether the cell-type specific SAM biosynthesis status is present and how it contributes to germline are not elusive.

Repression of retrotransposons is a crucial matter for the germline. Since they contain their cisregulatory elements for their transcription and their mobilization is caused by a “copy and paste” manner, the transposition of retrotransposons causes deleterious outcomes to the germline genome by either adding new regulating elements or destroying gene sequences. Since the nature of retrotransposons could cause catastrophic consequences for the host germline genome, how transposon expression is repressed in the germline is crucial.

In addition to their harmful effects, some retrotransposons were utilized in host cellular functions. One example is the retrotransposon’s function in telomere maintenance. In *Drosophila*, which lacks telomerase, telomere maintenance depends on the telomere-specific retrotransposons. *Drosophila* telomere retrotransposons consist of three non-Long Terminal Repeat (LTR) retrotransposons, *HeT-A*, *TART* and *THARE* (6). These elements are repeatedly located at the ends of chromosomes to make telomere repeats. Among those, *HeT-A* is the most abundant element and is well studied for its regulation (7, 8). *HeT-A* is approximately 6 kbp in length and contains a single open reading frame encoding Gag protein required for its localization to chromosome ends (9–11). Within the last 600bp of its 3’Untranslated-region (UTR), the promoter regulating transcription of adjacent downstream *HeT-A* element is located (12, 13). Once the promoter is activated, the transcribed RNA intermediates form a Ribonucleoprotein (RNP) complex with the Gag protein translated from itself. The formed RNP complex locates to chromosome ends mediated by Gag protein function, where *HeT-A* RNA is reverse-transcribed and integrated into telomere repeat (9). Although *HeT-A* and other telomere retrotransposons have useful functions to host chromosome protection, precise regulation of those elements is essential; it has been shown that excess expression of *HeT-A* causes insertions of it outside of the telomere (11, 14). Those evidences emphasize the importance of regulating retrotransposon expression in the germline.

In the *Drosophila* germline, retrotransposons are mainly regulated by a PIWI-interacting RNA (piRNA) -mediated repressive mechanism, which induces the formation of heterochromatin and the degradation of RNA intermediates (9, 15–17). In addition, a recent study showed that N^6^-methyladenine (6mA) DNA modification is involved in the positive regulation of retrotransposon expression (18). 6mA DNA modification is a common DNA modification of bacterial genome (19) and is also found in higher eukaryotic genome, including worms and flies (18, 20). Although the molecular function is not well understood, it has been shown that enrichment of 6mA up-regulates retrotransposon expression. Even though the molecular mechanism regulating retrotransposons during the *Drosophila* germline has been intensively studied, how germline metabolic status contributes to its regulation is unknown.

In this study, we provide the first evidence showing that the *Drosophila* germline in gametogenesis has a unique repressive SAM biosynthesis status. Our data indicates that repressive SAM biosynthesis status in the germline contributes to repression of retrotransposons, especially the telomeric element, *HeT-A*. Through functional analysis, we found that increasing SAM enhances the 6mA modification in the *HeT-A* promoter and its activity. Therefore, our study provides a model suggesting that germline unique SAM biosynthesis status is the new regulatory layer for controlling retrotransposon expression in the germline.

## Results

### SAM biosynthesis during gametogenesis

*Drosophila* gonads consist of two fundamentally different cell lineages, germline and somatic line. *Drosophila* ovaries consist of approximately 16 ovarioles, and at the anterior of each ovariole, ovarian somatic cells functioning as niche cells for germline stem cells (GSCs) are located (Figure 1B). Those niche cells (Terminal filament and cap cells) maintain associated GSCs (21). The differentiating daughter cells (Cystoblasts: CB) provided by GSC asymmetric division undergo 4 times mitosis with incomplete cytokinesis to be germline cysts. Germline cysts are then encapsulated with somatic follicle cells to form egg chambers (Figure 1B). To evaluate SAM biosynthetic status in ovarian cells, we generated an anti-Sam-S antibody to observe Sam-S protein expression. Surprisingly, we found that Sam-S expression is not observed in all germline cells, including GSCs and developing germline cysts, though Sam-S expression is stably observed in most of the ovarian somatic cells except the niche cells (Figures 1C, C’). We also observed Sam-S expression in a later stage of oogenesis and found it contentiously low in the germline, indicating that SAM biosynthesis is suppressed in the germline during oogenesis (Figures 1D, D’). Then we investigated whether the suppressed status of SAM biosynthesis also occurs in male germline. Similar to the ovarian case, somatic niche cells (hub cells) are located in the anterior region of the testis. Hub cells maintain associated GSCs and somatic cyst stem cells (CSCs). GSCs provide differentiating daughter cells, which form germline cysts; those germline cysts are encapsulated with cyst cells provided from CSCs to form cysts. Consistent with the ovarian case, we found that Sam-S expression is low in the germline during spermatogenesis (Figures 1E, E’). Conversely, the somatic expression of Sam-S is high except for the niche cells (Figures 1E, E’). Taken together, we conclude that the germline in gametogenesis has a suppressed status for SAM biosynthesis.

### The role of germline unique SAM suppression during oogenesis

To investigate the role of germline-unique suppressive SAM biosynthesis status, we overexpressed *Sam-S* in the germline by using germline-specific Gal4-driver, *nanos* (*nos*)-Gal4 combined with UASp-*Sam-S* (*nos* > *Sam-S*) (3). Overexpression of *Sam-S* certainly caused the expression of Sam-S protein in the germline, including GSCs (Figures 1F, F’). We also confirmed that the overexpression of *Sam-S* in the germline enhances SAM biosynthesis in ovaries by measuring amounts of SAM and its related metabolites. Our results showed that SAM and its catabolized compound, SAH, were significantly increased by overexpression of *Sam-S* in the germline (Figures 1H, I), with a slight decrease of its precursor compound, Met (Figure 1G). Those results confirm that SAM biosynthesis in the germline depends on the expression of Sam-S and suggest that generated SAM is catalyzed as a methyl base donor.

Previously, we reported that the overexpression of *Sam-S* caused excess germline cell death in the region where the DNA double-strand break check point works (3). This phenomenon resembles the effects of de-repression of retrotransposons caused by such as disruption of piRNA biosynthesis (14, 22). Therefore, we hypothesized that the overexpression of *Sam-S* results in the expression of retrotransposons in the germline. To test this, we performed quantitative reverse transcription PCR (qRT-PCR) on several retrotransposons, including two major telomeric elements, *HeT-A* and *TART*. We found that expressions of 10 out of 14 retrotransposons tested were elevated in ovaries with *Sam-S* overexpression in the germline (Figure 2). Among them, the *HeT-A* was the most affected retrotransposon, which had an approximate 7-fold increase in its expression (Figure 2). Thus, our results indicate that the Sam-S expression level in the germline is one of the regulators for retrotransposon expression, and low-level expression of Sam-S contributes to the repression of retrotransposon expression in the germline.

**Figure 2.**
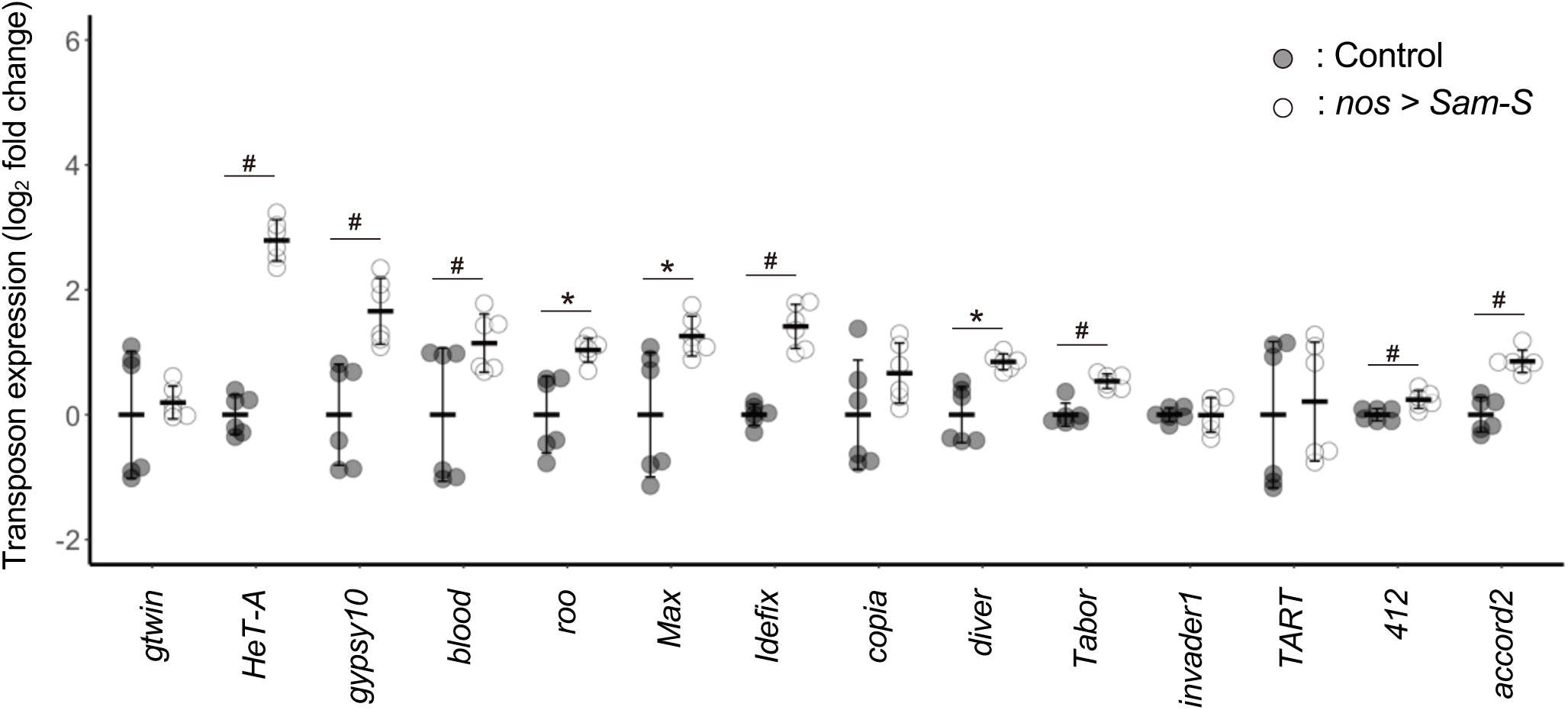
Retrotransposon expression in ovaries with SAM-increase germline. Comparison of expression of indicated retrotransposons is shown. Gray and white indicate the retrotransposon expression in control or *nos* > *Sam-S* ovaries, respectively. # : p <0.05 (Student’s t-test). *: p<0.05 (Welch’s t-test).

### Mechanisms for retrotransposon expression by SAM biosynthesis status

Next, we tried to address how SAM biosynthesis in the germline regulates retrotransposon expression. In general, piRNA-dependent transcriptional and post-transcriptional regulation is the major regulatory mechanism for retrotransposon expression. Thus, we first checked if the *Sam-S* overexpression in the germline alters the piRNA synthesis. In the *Drosophila* germline, 23- to 30- nt piRNAs are generated from long precursor RNAs with the activity of the PIWI clade of Argonaute proteins, Piwi, Aubergene (Aub), and Ago3 (14). This piRNA synthesis occurs in the subcellular region called Nuage, which is located peripherally to the nuclear membrane. We checked if the overexpression of *Sam-S* in the germline affects the expression and localization of Argonaute proteins and piRNA synthesis. Our results showed that the expression and sub-cellular localization of investigated Argonaute proteins were not affected by *Sam-S* overexpression (Supp. Figures 1A-C, Supp. Figure 2) and also showed no alternation to the amount and size of synthesized piRNA (Supp. Figure 1D). Thus, excess expression of retrotransposons in *Sam-S* overexpressed germline is not assumed to be caused by change of piRNA synthesis.

Next, we sought the other regulatory points affected by the SAM increase. A recent study has shown that the DNA 6mA methylation modification positively regulates the expression of retrotransposons (18). Therefore, we investigated whether the increase of retrotransposon expressions in SAM-increased germline is caused by the increase of 6mA modification. To this end, we performed methylated-DNA immunoprecipitation sequencing (MeDIP-Seq) analysis using an anti-6mA antibody.

Since a recent study argues that the detection of 6mA in animal genomes is caused by contaminations of RNA or the endosymbiotic bacterial genomes (23), we established the genome extraction procedure in which those phenomena could occur less. First, we determined the condition in which the RNA contamination less occurs by performing consecutive RNase treatments. Our results suggested that two treatments of RNase are enough to remove almost all of the RNA from extracted DNA samples (Supp. Figure 3). Next, we checked whether extracted genomic DNA samples contain the endosymbiotic bacterial or food-derived yeast genomes by PCR (24, 25). Our results showed that genomic DNA extracted from ovaries did not contain contaminated genomes from those species, even when whole-body genomic samples contain those genomes (Supp. Figure 4). From those results, we conclude that contamination of those materials occurs less in our ovarian genomic DNA samples.

With the above samples, we performed MeDIP-Seq on the genomes from the control and *Sam-S* overexpressed ovaries. We mapped the obtained reads (Supp. Table 2) to the sequences of retrotransposons; those expressions were enhanced by *Sam-S* overexpression in the germline. We found significant enhancement of 6mA modification in the 5’UTR and 3’UTR of *HeT-A* in *Sam-S* overexpressed germline (Figure 3A, Supp. Figure 5A). Of note, the 6mA enhancement by *Sam-S* overexpression occurred in the *HeT-A* promoter in its 3’UTR, suggesting that Sam-S dependent SAM increase regulates *HeT-A* promoter activity (Figure 3A). In contrast to *HeT-A*, enhancement of 6mA modification by *Sam-S* overexpression did not occur in the other retrotransposons (Figure 3B, Supp. Figures 5B, C, Supp. Figures 6, 7), suggesting that the increase of retrotransposon expression in *Sam-S* overexpressed germline is not caused by 6mA enhancement, and *HeT-A* is under the control of a unique regulatory mechanism dependent on SAM biosynthesis and 6mA modification. We checked if the obtained reads were mapped to the symbiotic bacterial genomes and confirmed that our reads merely mapped to those (Supp. Table 2).

**Figure 3.**
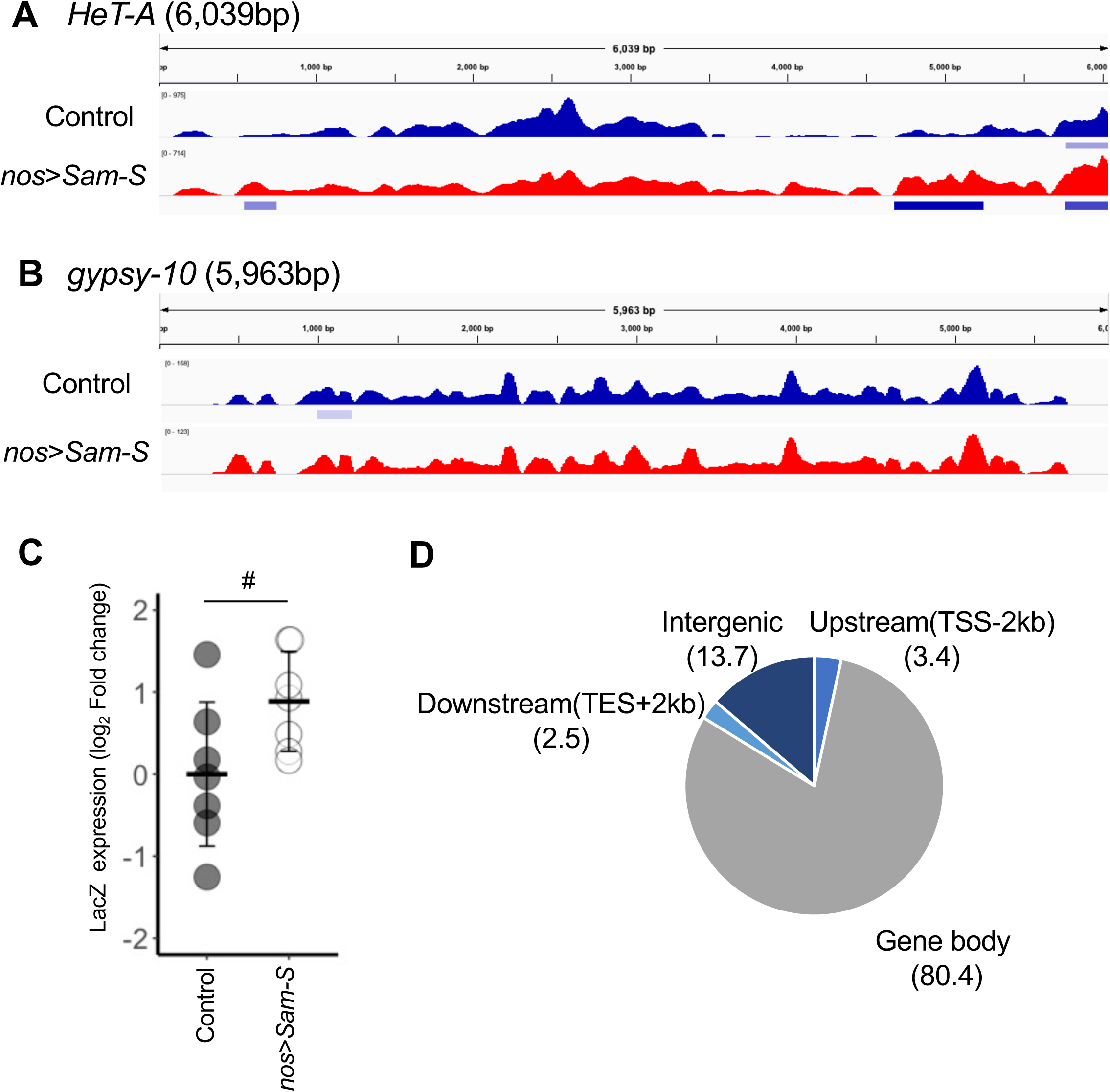
Effects of SAM increase to 6mA modification and *HeT-A* promoter activity. (A, B) 6mA enrichment to *HeT-A* (A) and *gypsy-10* (B) are shown. Blue indicates the 6mA enrichment from control samples, and red indicates the results from *nos* > *Sam-S* samples. The bars underneath each panel indicate the detected peaks, and the darker bar color indicates a higher MACS score. (C) Comparison of reporter activity for *HeT-A* promoter is shown. Gray and white indicate reporter (LacZ) expression in control or *nos* > *Sam-S* ovaries, respectively. #: p < 0.05 (Student’s test). (D) The genomic location of unique peaks detected in *nos* > *Sam-S* ovaries. The numbers in the brackets represent each category’s percentage (%).

We next investigate if an increase of SAM biosynthesis affects the *HeT-A* promoter activity. To test this, we utilized reporter to *HeT-A* promoter activity, in which LacZ coding sequence is under the control of *HeT-A* promoter sequence (13, 26). We performed qRT-PCR to investigate LacZ expression in control and *Sam-S* overexpressed ovaries. We found a significant increase of LacZ expression in *Sam-S* overexpressed ovaries (Figure 3C), indicating that *HeT-A* promoter activity is controlled by SAM biosynthesis.

### Effects of SAM biosynthesis on germline gene expression

We next investigated the effects of SAM-dependent increase of 6mA modification to gene expression outside of retrotransposons. We mapped the MeDIP-Seq reads to the whole fly genome and extracted the peaks observed only in SAM-increased condition. We detected 925 peaks uniquely observed in the *Sam-S* overexpressed condition. We found that most of the unique peaks are located upstream of genes or gene bodies (775 peaks out of 925 total unique peaks, Figure 3D). Thus, we speculated that SAM-dependent 6mA enhancement affects gene expression. To test this, we performed RNA-sequence (RNA-Seq) analysis to see the effects of SAM increase to the total gene expression in the germline. Our RNA-Seq analysis detected 122 differentially expressed genes from 17457 genes totally detected (30 genes: up-regulated, 92 genes: down-regulated) (Figure 4A). *Sam-S* was one of the most affected genes, confirming that our RNA-Seq analysis works properly (Figure 4A). Although our RNA-Seq analysis performed well, we detected a relatively small population of genes affected by SAM increase. In addition, we found there are few structural genes involved in germline development, except female *sterile*(*1*)*K10* encoding a nuclear protein required for proper oocyte-polarity formation (27) (Figure 4A). Also, gene ontology (GO) enrichment analysis for differentially expressed genes (DEGs) showed no GO related to germline development (Figure 4B), suggesting that increasing the amount of SAM in the germline did not have obvious effects on the proper progression of germline development. Finally, we investigated the relation between genes with 6mA peak(s) and DEGs uniquely observed in the SAM increase condition. We found that most of the genes with altered 6mA modification did not merge with the DEGs (Figure 4 C). Those results suggest that SAM biosynthesis-mediated 6mA modification is not a major regulator of the germline gene expression during oogenesis and emphasize its unique regulatory role in *HeT-A* transcriptional regulation.

**Figure 4.**
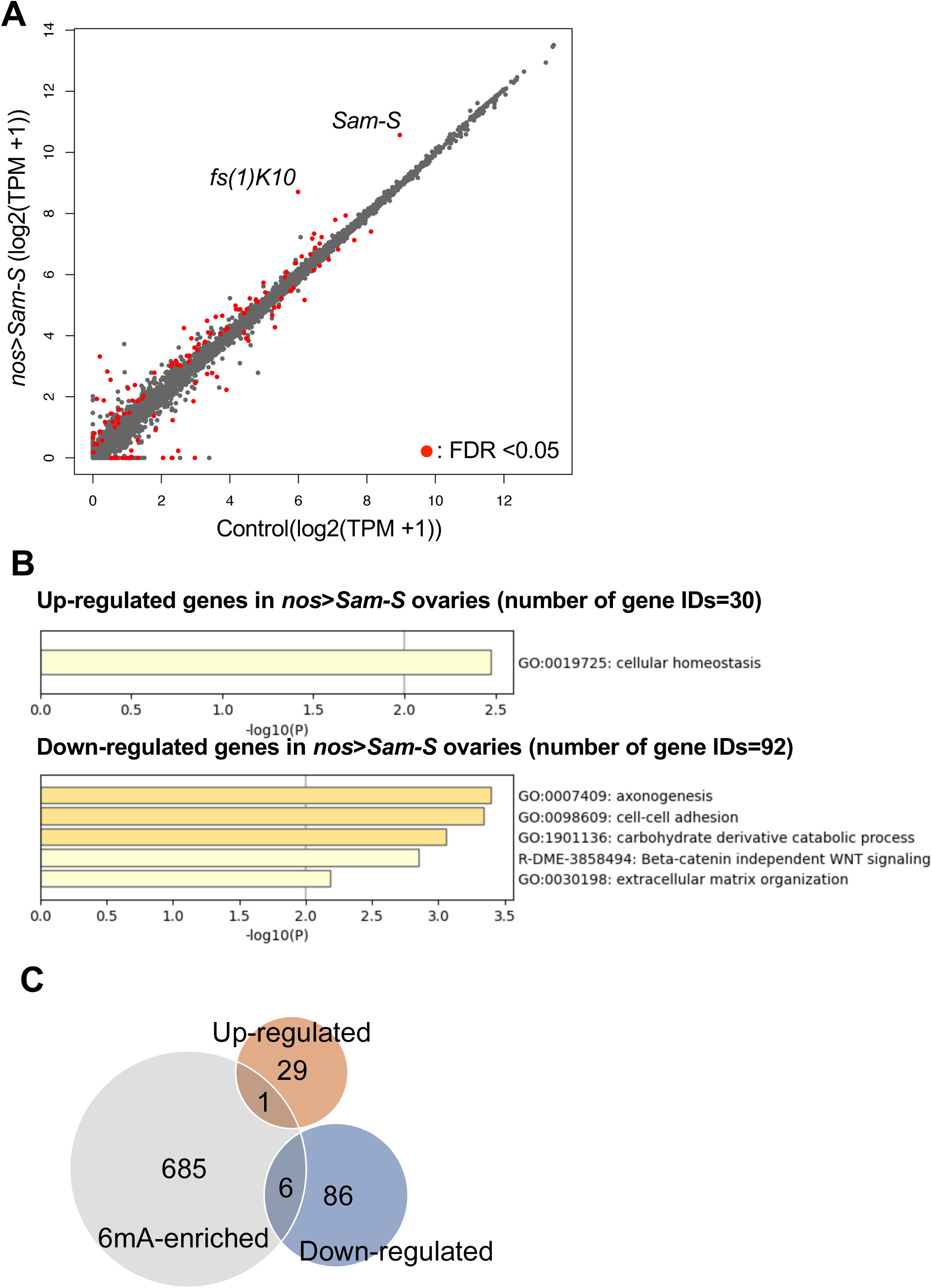
Effects of SAM increase to gene expression in ovaries. (A) Comparison of RNA-Seq. data from control and *nos* > *Sam-S* ovaries. The red dots indicate the differentially expressed genes determined with False Discovery Rate (FDR). (B) GO enrichment analysis for genes up-regulated (top) or down-regulated (bottom) in *nos* > *Sam-S* ovaries. GO enrichment analysis was performed with the Metascape web tool. (C) The relation between the genes with 6mA peak(s) uniquely observed in *nos* > *Sam-S* ovaries (Represented as “6mA-enriched”) and the genes those expression was affected in *nos* > *Sam-S* ovaries (Represented as “Up-regulated” or “Down-regulated”). The numbers represent the number of the genes in each category.

## Discussion

SAM is the fundamental metabolite for cellular function, providing methyl-moiety for methylation reactions. Since SAM is crucial for cellular function and is thought to be synthesized in principally all cell types (28), the cell-type-specific status for SAM biosynthesis remained elusive. In this study, we first reported that the *Drosophila* germline in gametogenesis has a repressive status for SAM biosynthesis by observing Sam-S, a single SAM biosynthetic enzyme encoded in the *Drosophila* genome. We also firstly revealed that this repressive status for SAM biosynthesis in the germline contributes to the repression of retrotransposons. Thus, our finding provides a new mechanistic layer for inhibiting retrotransposon expression in the germline. Since this mechanism depends on a simple mechanism, suppression of Sam-S in the germline, and it is known that some eukaryotes, such as plants, lack piRNA synthetic pathway (29), it is possible that our found SAM-biosynthesis-dependent suppression of retrotransposons could be the ancestor mechanism regulating retrotransposons in the germline.

We previously reported that the age-dependent increase of *Sam-S* expression in the germline is the cause of the age-dependent decline of egg production ability without particular molecular insight regulating this phenomenon (3). Given that DNA damage is thought of as one of the causes of cellular/ tissue aging, it is worthwhile to note that the SAM-dependent increase of retrotransposon expression in the germline is a possible candidate for germline aging. In our gene expression analysis, we found some GOs related to cell-cell interactions are altered in SAM increase condition, and one of the age-related disorders during *Drosophila* oogenesis appeared in egg chamber formation (3), it is formally possible that SAM-dependent germline aging is caused by the alternation of those gene expressions. Our research is the first report showing the effects of SAM synthesis on germline gene expression in *Drosophila*.

As one of the molecular mechanisms regulating retrotransposon expression underneath the alternation of SAM biosynthesis, we focused on the possibility that the transcription of retrotransposon is regulated via 6mA modification since previous work showed that the accumulation of 6mA modification has a positive correlation to retrotransposon expression, including *HeT-A* (18). Our results showed that increased SAM causes significant enrichment of 6mA in the *HeT-A* promoter region. Moreover, we showed that *HeT-A* promoter activity was enhanced in an SAM-increased germline. We did not observe any correlation between 6mA accumulation and expression in most retrotransposons; those expressions are up-regulated in an SAM-increased condition. Thus, our results suggest that *HeT-A* transcription is controlled by a unique regulatory mechanism dependent on SAM-mediated regulation of 6mA modification. Although we could not exclude the possibility that some other methylation-dependent transcriptional regulatory mechanism, such as methylation on Lysine 4 of Histone H3, the general epigenetic modification involved in transcriptional activation, is also involved in this process, the strong correlation between the specific accumulation of 6mA on the promoter region and the enhancement of its activity suggests that 6mA regulates the promoter activity of *HeT-A*. How 6mA modification regulates *HeT-A* promoter activity and whether the other methylation modification is involved in its regulation are questions to be asked in future studies.

A recent study argues that the presence of 6mA DNA modification in eukaryotic genome comes from the contamination of RNA and/or the endosymbiotic bacterial genome, which is known to have 6mA modification (23). Although this could be the case for the genomic samples from whole fly bodies with which we detected contamination of symbiotic bacteria or food-derived yeasts, we did not detect that contamination in our genomic sample extracted from dissected ovaries. Also, our consecutive RNase treatments showed that two treatments of RNase are enough to remove almost all of the RNA from our genomic samples. Thus, our study reinforces the presence of 6mA modification, at least in the *Drosophila* ovarian genome. At the same time, our study showed that the contribution of SAM-mediated regulation of 6mA modification to gene expression is quite limited. Thus, although 6mA modification is present in the *Drosophila* genome, solo enhancement of 6mA modification does not affect the expression of structural genes and retrotransposons except *HeT-A*.

SAM has been thought to be an upstream regulator of gene expression. We firstly show the effects of a solo SAM increase on gene expression in the germline during gametogenesis. Although it affects some gene expression, the effects are relatively minor, especially for genes involved in germline development. Conversely, most retrotransposons we tested were affected by their expression in SAM increase conditions. Those results provide insight into the differences in regulatory rigidness for gene expression between genes involved in developmental processes and the genetic material added from outsides; regulation of structural genes is strongly controlled by the molecular machinery, such as “writer” enzymes of epigenetic modification, conversely, retro-element regulation has less rigidity and is easily affected by the amount of donor material. Those less rigid regulations on retroelements may provide the relatively sensitive response of their expression to cellular physiological conditions (30).

In this study, we provide the model that the germline unique repressive status of SAM biosynthesis contributes to the regulation of retrotransposon expression, including the telomeric element, *HeT-A*. This mechanism depends on the function of unidentified factor(s) that regulate the suppression of Sam-S in the germline. By identifying the factor(s) regulating Sam-S expression in future studies, we could understand how germline unique metabolic status is established and how it contributes to regulating germline genome integrity and aging. Our found phenomena could be the important first step in understanding the mechanism interconnecting germline unique metabolic status and its contribution to the continuity of life.

## Materials & Methods

### Fly strains

Flies were maintained on standard *Drosophila* medium at 25 ℃. germline-specific Gal4 driver *nanos*-Gal4::VP16 (*nos*-Gal4, Gift from D. Van Doren)(31) was crossed to UASp-*Sam-S* (3) or M[3xP3-RFP. attP’]ZH-86Fa (#24486 from BDSC, control for pUASp-*Sam-S*). *HeT-A*-LacZ (H111) strain was kindly gift from Dr. Kalmykova (26).

### Immunohistochemistry

Antibody staining was performed according to standard procedures (32). Antibody staining was performed according to standard procedures (32). The following primary antibodies were used: mouse anti-1B1 antibody [1:5, Developmental Studies Hybridoma Bank (DSHB)], mouse anti-FasIII antibody (1:5, DSHB), chick anti-Vasa antibody (1:500, lab stock), and rabbit anti-Sam-S antibody (1:500, this work). Stained ovaries were mounted in Vectasheild (Vector Laboratories, U.S.A.) and imaged with a confocal microscope (TCS SP5, Leica Microsystems, Germany).

### Generation of antibody

Anti-Sam-S antibody was generated in rabbit via immunization with a synthetic peptide corresponding to amino acids 95-108 (RETVQHIGYDDSSKG) of Sam-S-PA (MBL, Japan).

### UPLC-MS/MS analysis

Detailed procedures for the extraction of metabolites and UPLC-MS/MS analysis were described previously(3). In roughly, ovaries metabolites were extracted from 20 female flies in 50% methanol. After the evaporation and filtration, samples were diluted with equal volumes of either 50 mM Tris-HCl (pH 8.8) with 100 µM DTT. Measurement of metabolites was performed using ultra-high-performance liquid chromatography equipped with tandem mass spectrometry (UPLC-MS/MS) (SHIMADZU, Japan).

### q-PCR

q-RT-PCR was performed as previously described (3). RNAs extracted from fly ovaries by Trizol (Thermo Fisher Scientific, U.A.S.) were reverse-transcribed with Super Script III Reverse Transcriptase (Thermo Fisher Scientific), and synthesized cDNA was used as a template for q-PCR with QuantiFast SYBR Green PCR Kit and Rotor-Gene Q thermal cycler (QIAGEN, Netherlands). Genomic q-PCR was performed the same way above, except the used template was prepared as described below. Primers used for qRT-PCR are listed in Supp. Table1.

### Preparation of ovarian genomic DNA

For ovarian genomic DNA (gDNA) extraction, ovaries from 20 female flies were used. Ovaries were dissected in PBS. Ovaries were flash-frozen in liquid nitrogen and crushed with a homogenizer (5000 rpm, 30 sec., 2 times: Minilys, M&S Instruments, Japan). Homogenized samples were treated with a Wizard genomic DNA purification kit (Promega, U.S.A.) following the manufacturer’s protocol. To avoid RNA contamination, we treated RNase A twice. For the times of RNase A treatments, we tested consecutive RNase A treatment. We determined the condition for removing almost all of the RNA by measuring the amount of nuclear acids with a microvolume spectrophotometer (DS-11, Denovix, U.S.A.).

### Detection of contaminated genomic DNA

The detection of contaminated genomic DNA from endosymbionts or food containing yeasts was done with PCR, as reported previously (24, 25). The genomic DNAs extracted from the whole body or ovaries were used to check the contamination. PCR was performed with PrimeSTAR Max DNA polymerase (TAKARA, Japan) with the primers listed in Supp. Table1. Annealing temperature for each PCR reaction is below.

Mitochondria and *Wolbachia pipentis*: 52℃ Gut bacteria: 65℃, Yeasts: 50℃.

Contamination of gDNAs of endosymbionts or food containing yeasts was also checked by sequencing (Supp. Table 2).

### MeDIP-Seq

Extracted ovarian gDNA was sonicated in a Covaris Focused-ultrasonicator S220 (Peak Incident Power: 175W, Duty factor:10%, Cycle per burst:200, Time: 180s, Covaris LLC, U. S. A.) to be in approximate 200 bp size fragments. After the size selection by AMpure XP beads (Beckman Coulter Inc., U.S.A.), 1µg of collected fragments from control or *Sam-S*-overexpressed ovaries were used for end-repair and adaptor ligation employing the NEB Next Ultra DNA Library Prep Kit for Illumina (E7370, New England Biolabs, U.S.A.). The fragments were denatured and processed for immunoprecipitation. After the removal of 10% input, the lest of fragments were incubated with 5µg of anti-6mA antibody (202-003, Synaptic Systems, Germany) in Immunoprecipitation buffer (50mM Tris-HCl pH8.0, 150mM NaCl, 0.1% TritonX-100) at 4℃ for overnight. The fragments bound with antibodies were collected with a mixture of protein A or G magnet beads (TAMAGAWA SEIKI, Japan) pre-treated with normal rabbit IgG (12-370, Merck Millipore, U.S.A.). The fragments were then purified by Proteinase K treatment followed by Phenol/Chloroform/Isoamyl alcohol precipitation. After the PCR amplification with sample-specific barcodes (Input sample: 12 cycles, IP samples: 15 cycles), the libraries were sequenced with NextSeq500 Desktop Sequencer (Illumina, U.S.A.). For data analysis, treated reads were mapped to each retrotransposon sequence or *Drosophila* genome (dmel_6.45) with bowtie2 (2.3.2), and peak detection was done with MACS (2.1.1.20160309) software. Input reads were used to normalize the peak detection. To extract the unique peaks in the *nos* > *Sam-S* condition, first, we extracted the common peaks detected in *nos* > *Sam-S* samples and subtracted the peaks detected in control samples. Then, the nearest gene regions from extracted peaks were detected by the ChIPpeakAnno package from Bioconductor (https://bioconductor.org/.).

### RNA-Seq

The extraction of RNA is performed as described above. Library preparation was performed with TruSeq Stranded mRNA Library kit (Illumima) following manufacturer’s instruction. High-throughput sequencing was performed with NextSeq500 Desktop Sequencer (Illumina). The obtained reads were treated with trim-galore and mapped to the *Drosophila* genome (dmel_r6.27) using STAR (2.7.1a). The expression of genes was calculated using the DESeq2 package from Bioconductor.

### Northern blot analysis

Northern blot was performed as previously described (33). Total RNA was isolated from the ovaries of 100 female flies by ISOGEN (Fujifilm, Japan). 5 µg of total RNA was resolved by electrophoresis. Probes used for *roo* and U6 snRNA were 5’- TCGACTCAGTGGCACAATAAAT-3’ and 5’-GGGCCATGCTAATCTTCTCTGTA-3’, respectively. The DNA oligos were labeled with T4 polynucleotide kinase in the presence of [γ-^32^P] ATP.

### Western blot analysis

Western blot was performed as previously described (34). Proteins in lysates were separated by SDS-polyacrylamide gel electrophoresis and transferred to a 0.45µm nitrocellulose membrane (Cytiva). The primary antibodies used were; mouse anti-Aub antibody (4D10, 10003Ab-1, NIG)(35), mouse anti-Piwi antibody (P4D2) (36) and anti-Tubulin antibody (E7, DSHB). Then, the membrane was treated with an HRP-conjugated secondary antibody (0855558, MP Biomedicals, CA, U.S.A.) for 30 min. at room temperature. The signal was developed with ECL Prime Western Blotting Detection Reagent (Cytiva) and visualized by a C-DiGit Blot Scanner (LI-COR Biotechnology, U.S.A.).

## Data availability

The sequencing data have been deposited in the Gene Expression Omnibus (GEO) under the accession numbers GSE294101 and GSE294178.

## Acknowledgement

We thank the Bloomington Drosophila Stock Center and Developmental Studies Hybridoma Bank for providing us with the materials. This work was supported in part by Grants-in-Aid for Scientific Research from Japan Society for Promotion of Science (JSPS) (KAKENHI Grant Number: 26116730, 18K06240 and 23K05777 to Y.H., 21H04774 and 24H00567 to M.M., 24H02030, 18H05552 to Sa.Ko.) and the Japanese Science and Technology Agency (JST) PRESTO (Grant Number: JPMJPR24N4 to So.Ka.) and also by the program of Inter-University Research Network for High-Depth Omics, Institute of Molecular Embryology and Genetics (IMEG), Kumamoto University.

## Author contributions

Y.H., S.H., D.S., and So.Ka. designed the experiments. Y.H., S.H., So.Ka., K.S. and B.S. performed the experiments. Y.H., T.S., and M.S. analyzed the data. M.M., M.N., and Sa.Ko. supervised the project. Y.H. wrote the manuscript with support from S.H. and N.K. All authors discussed the results and commented on the manuscript.

---

**Supp. Figure 1.**
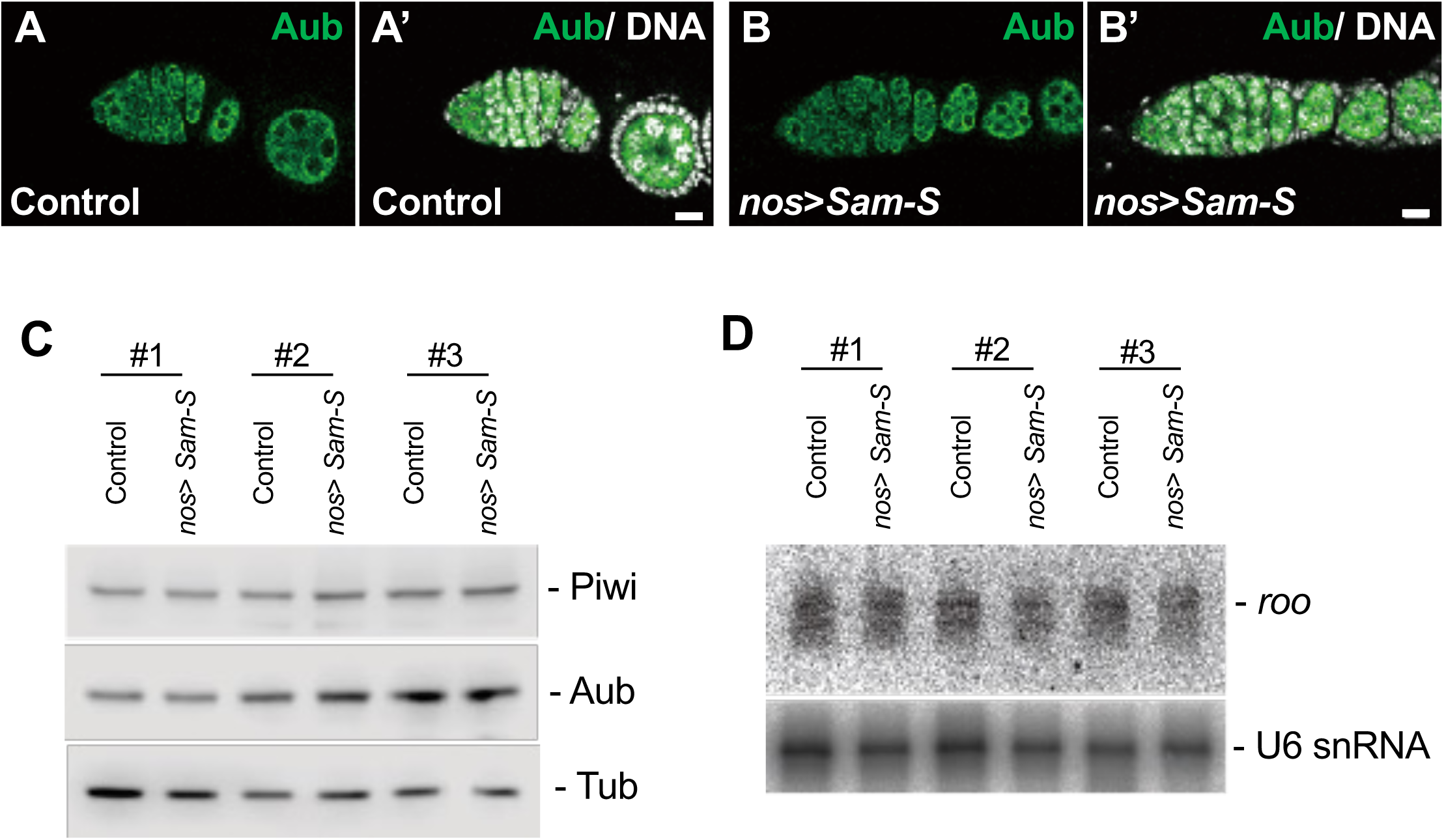
Effects of SAM increase to piRNA synthesis. (A-B’) Expression of Aub protein (green) in early oogenesis of control (A, A’) and *nos* > *Sam-S* females (B, B’) are shown. White signals in A’ and B’ indicate DNA. Expression and localization of Aub protein is not affected by germline *Sam-S* overexpession. (C) Western blot results of Piwi and Aub proteins are shown. There is no detectable difference between control and *nos* > *Sam-S* samples. Whole membrane images are shown in Supp. Figure 2. (D) Northern blot results of *roo* piRNA are shown. There is no detectable difference between control and *nos* > *Sam-S* samples. Bars in A’ and B’: 10µm.

**Supp. Figure 2.**
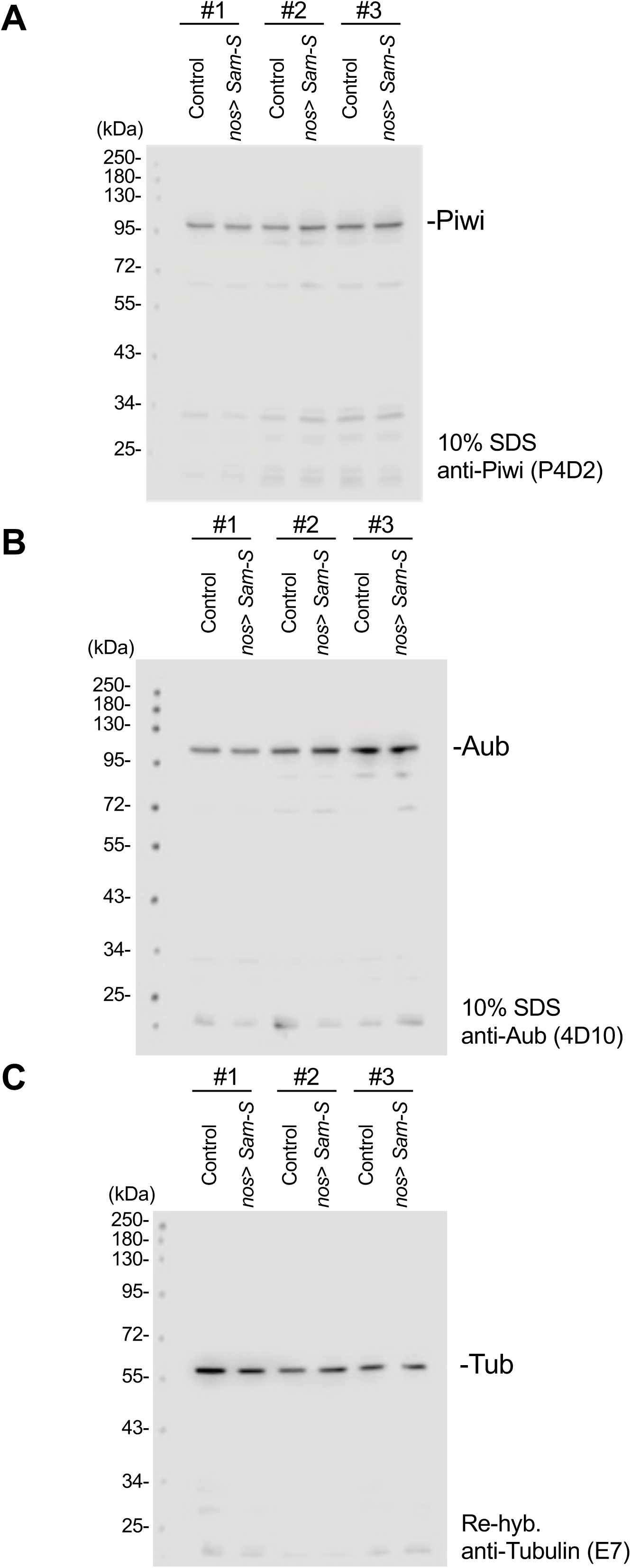
Whole membrane image of western blot results shown in supp. figure 1 C. Whole membrane image of western blot results for Piwi (A), Aub (B) and Tublin (C) are shown.

**Supp. Figure 3.**
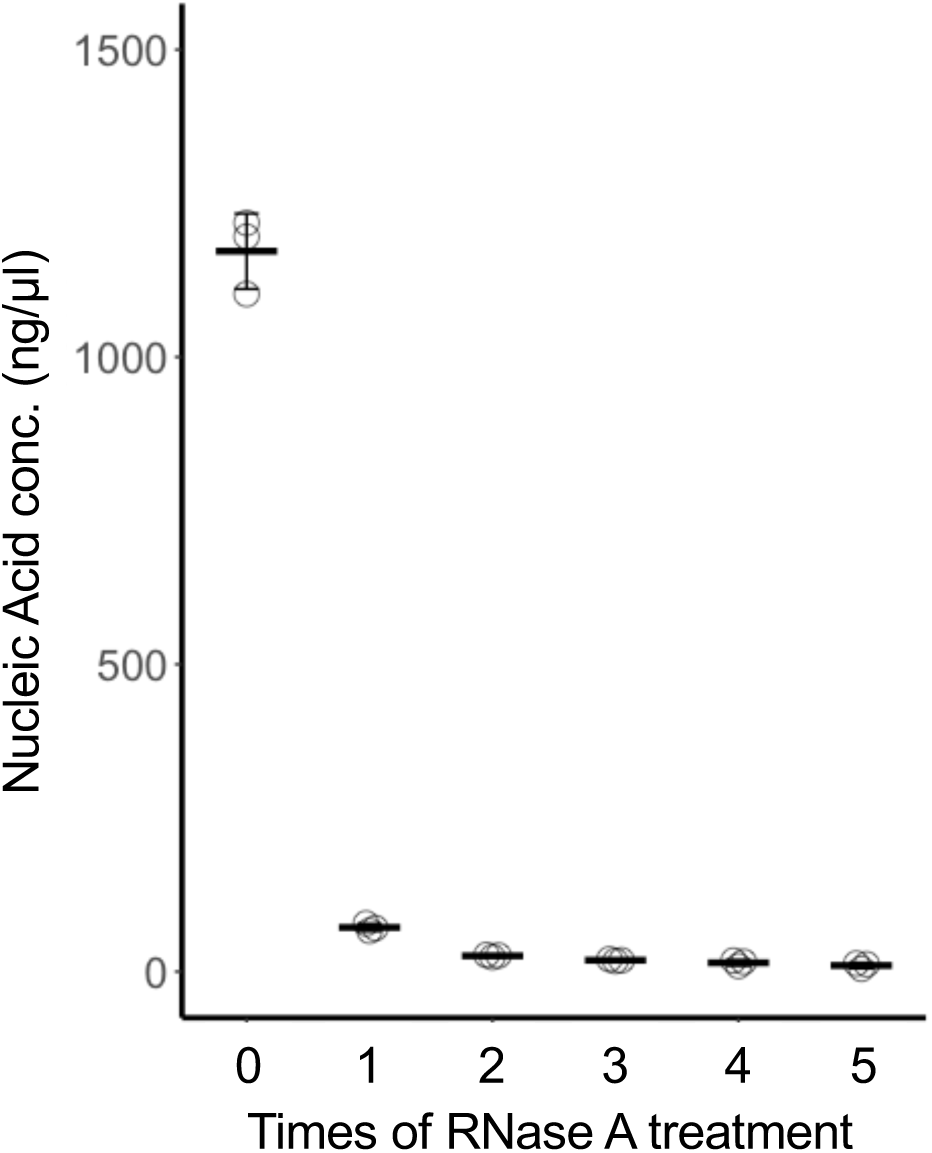
Effects of consecutive treatments of RNase A to the nucleic acid amount. The relation between the times of RNase A treatments and the amount of nucleic acid that resulted is shown. Three ovarian genome-extraction samples were consecutively treated with RNase.

**Supp. Figure 4.**
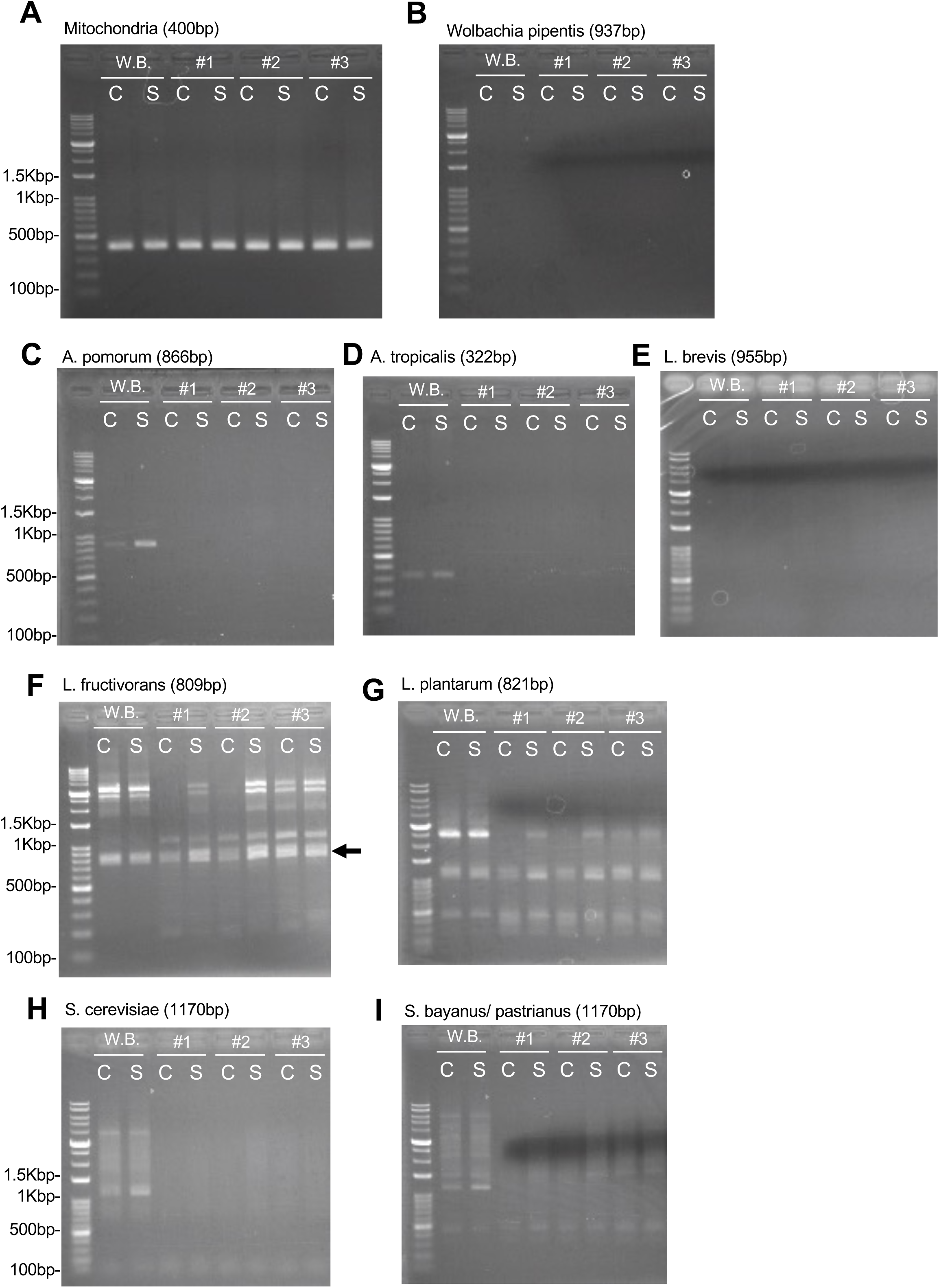
Investigation of genomic contamination of endosymbiont or yeasts. (A) Detection of mitochondrial genome as positive control for the following experiments. Genomic samples from the whole bodies (W.B.) and ovaries (#1-3) of control (C) or *nos* > *Sam-S* (S) females are PCR tested to investigate the presence of mitochondrial genome. (B-I) Same combination of samples were PCR tested to investigate the presence of *Wolbachia pipentis* (B), *Acetobacter pomorum* (C), *Acetobacter tropicalis* (D), *Lactobacillus brevis* (E), *Lactobacillus fructivorans* (F), *Lactobacillus plantarum* (G), *Saccharomyces cerevisiae* (H) and *Saccharomyces bayanus* or *pastrianus* (I). The expected product sizes are also indicated. In the case of *L. fructivorans* (F), we detected the PCR product having the similar size to expected size (arrow). To investigate if the PCR product was amplified from genome of *L. fructivorans*, we extracted the band and check with sequencing. We found that the product sequence is 95% matched with *Drosophila* chromosome 2R, thus it did not indicate the contamination of *L. fructivorans*.

**Supp. Figure 5.**
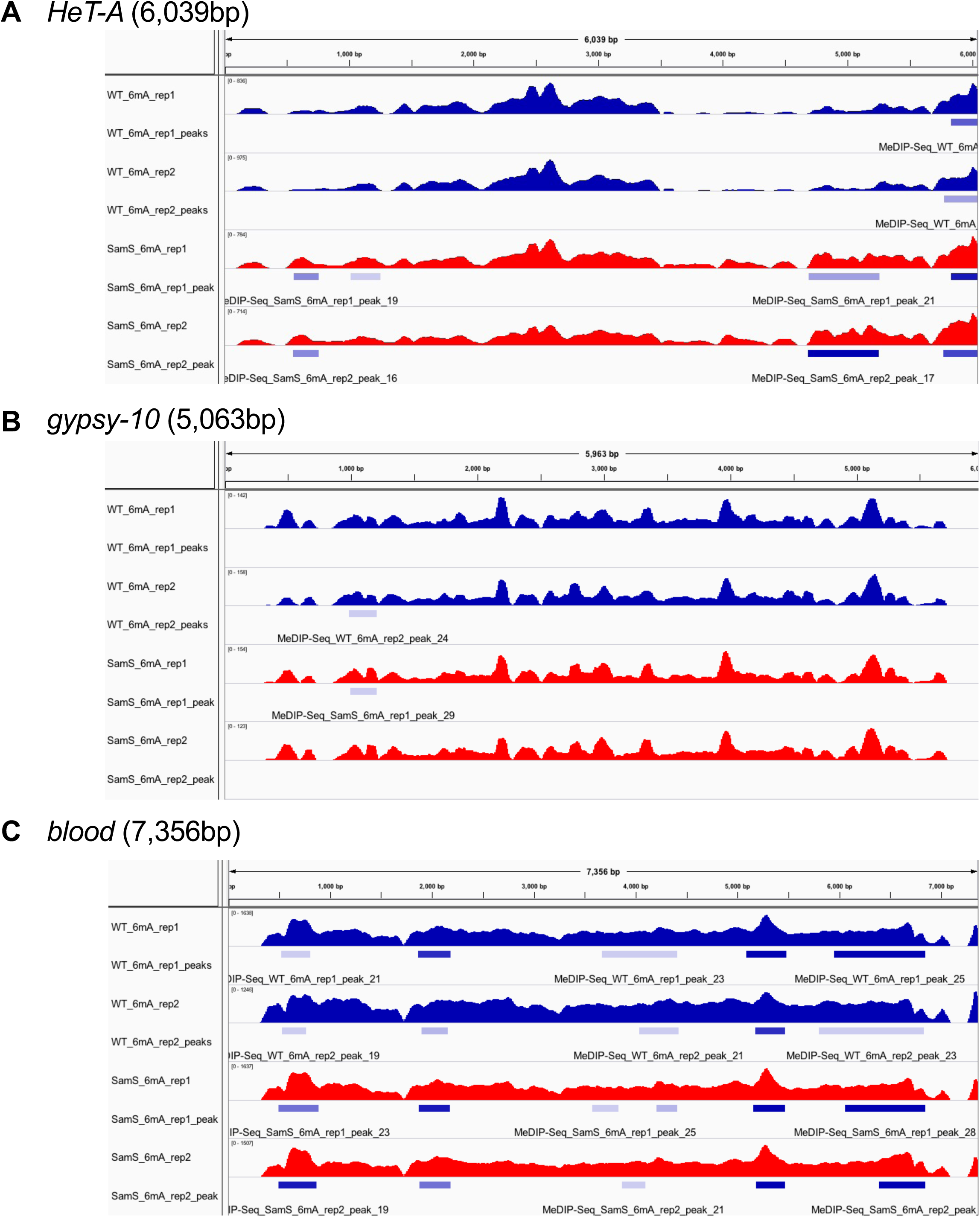
Enrichment of 6mA DNA modification to retrotransposon sequences (i). Enrichment of 6mA DNA modification in the *HeT-A* (A), *gypsy-10* (B), and *blood* (C) are shown. Duplicated ovarian genomic samples from control or *nos* > *Sam-S* females were investigated. The bars underneath the mapped reads indicate the detected peaks.

**Supp. Figure 6.**
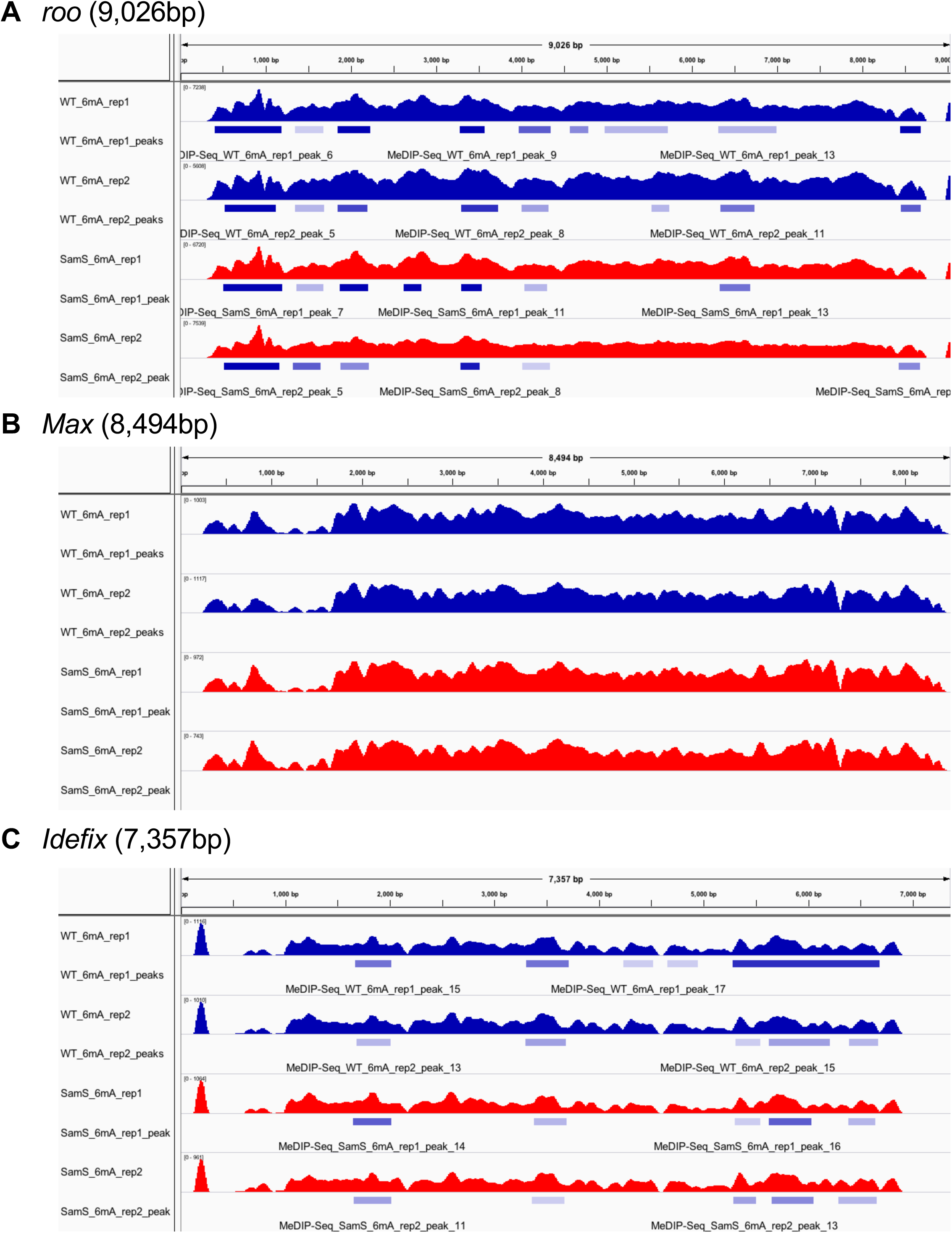
Enrichment of 6mA DNA modification to retrotransposon sequences (ii). Enrichment of 6mA DNA modification in the *roo* (A), *Max* (B), and *Idefix* (C) are shown. Duplicated ovarian genomic samples from control or *nos* > *Sam-S* females were investigated. The bars underneath the mapped reads indicate the detected peaks.

**Supp. Figure 7.**
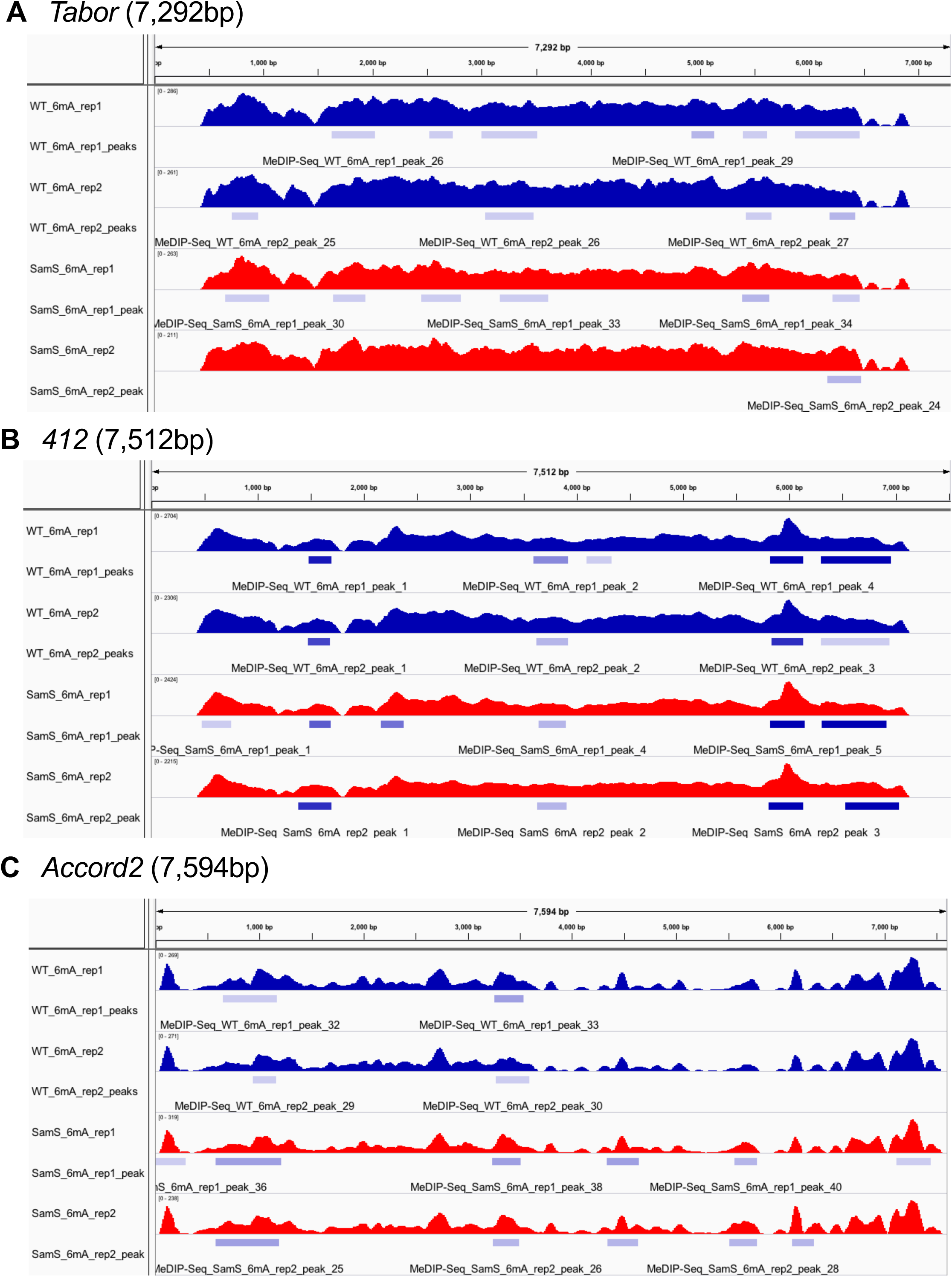
Enrichment of 6mA DNA modification to retrotransposon sequences (iii). Enrichment of 6mA DNA modification in the *Tabor* (A), *412* (B), and *Accord2* (C) are shown. Duplicated ovarian genomic samples from control or *nos* > *Sam-S* females were investigated. The bars underneath the mapped reads indicate the detected peaks.

**Supp. Table 1:**
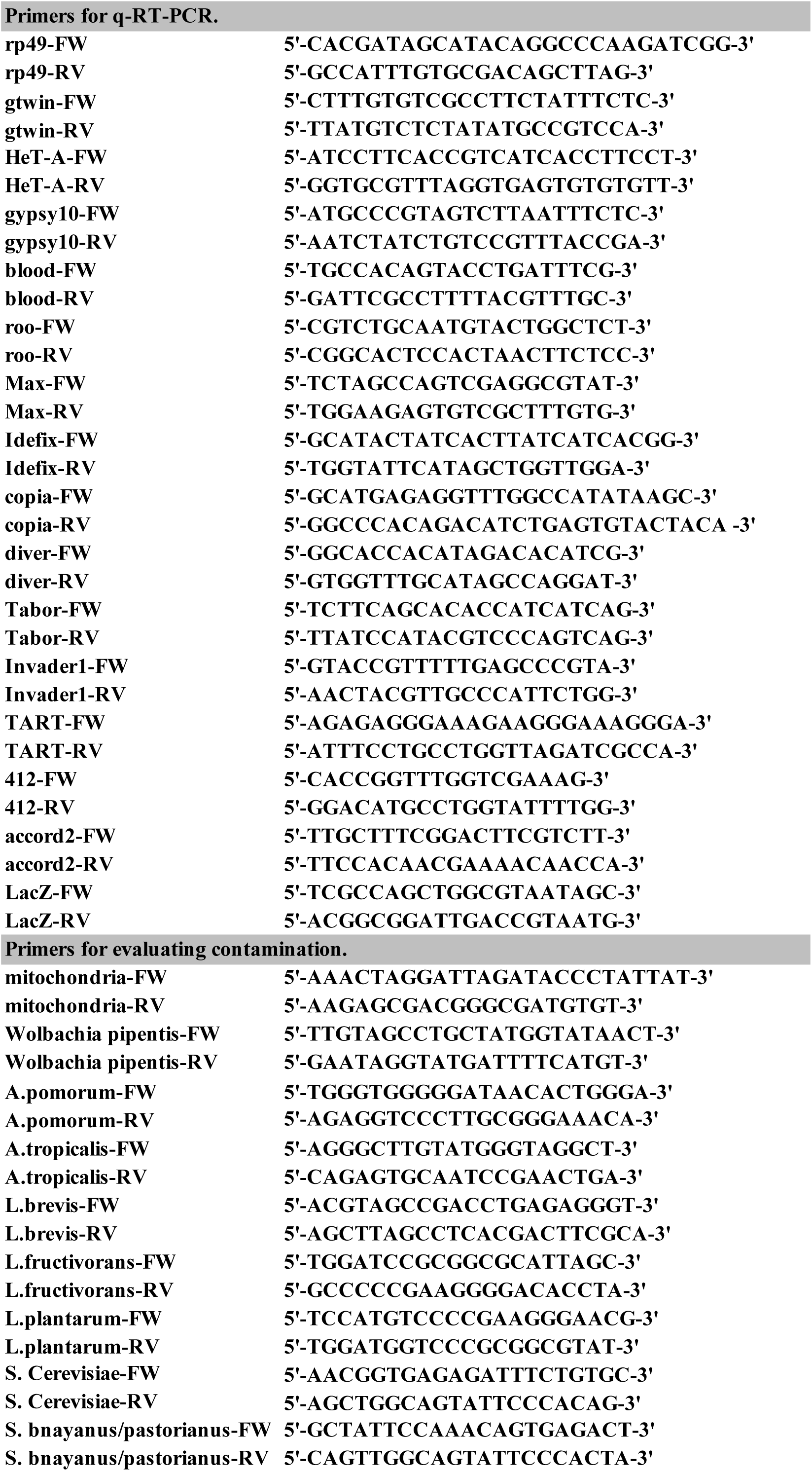
Primer list.

**Supp. Table 2:**
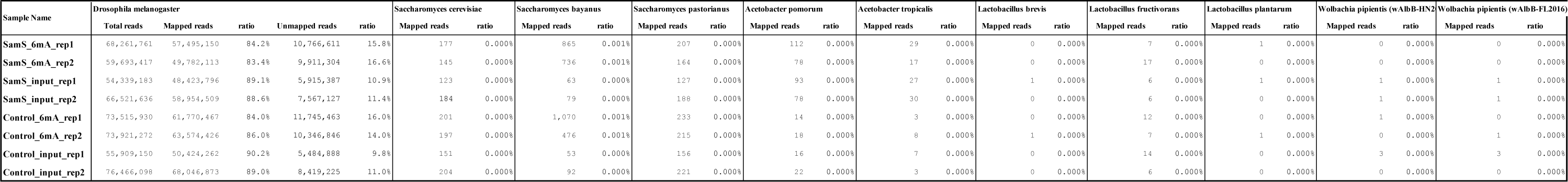
Mapping results of MeDIP-reads.

